# hmde: Hierarchical Methods for Differential Equations

**DOI:** 10.1101/2025.01.15.633280

**Authors:** Tess O’Brien, Fonti Kar, David Warton, Daniel Falster

## Abstract

Repeat observations of size are a common tool for understanding growth across taxa, and are used to estimate parameters for functions that describe growth rates. Recent advances in estimating differential equation parameters with a hierarchical longitudinal model have gone beyond available software. Custom implementation of such models is a barrier to use particularly for people who are not familiar with statistical programming. We introduce a new R package implementing a hierarchical Bayesian longitudinal model for repeat observation data with three growth models from ecological case studies. The package provides tools for model fitting and estimate extraction, example data, and case studies to demonstrate the use-case for each of the example models.

---

This paper describes hmde (https://github.com/traitecoevo/hmde), a new R package that fits a hierarchical model to estimate parameters of a differential equation (DE) in the presence of measurement error, where the parameters may vary randomly across subjects, following O’Brien et al. (2024). hmde implements a set of hierarchical Bayesian longitudinal models to fit DEs to repeat observation data and is an example of both a Bayesian method (Idier, 2013) and a mixed effects model. The package name stands for **h**ierarchical **m**ethods for **d**ifferential **e**quations. The model estimates DE parameters from repeated observations of the process over time. The motivating application for this package comes from ecology, where a common form of data is repeated observations (with error) of organism size, with the aim of estimating growth trajectories which may vary across individuals. The underlying statistical method was first used in O’Brien et al. (2024) to model tree growth, and we continue to focus on ecological applications in this paper with additional models and taxa. Existing methods such as non-linear mixed effects models have been used for tree growth with pair-wise difference data (de Miguel et al., 2013), and hierarchical models have been developed which fitted species-level trajectories (Her- ault et al., 2011), but the addition of the individual longitudinal structure required a new approach. This software allows for precise control over the hierarchical structure, differential equation solution, and estimation method. Three ordinary differential equation (ODE) models are provided: constant, a linear first-order ODE known as the von Bertalanffy model (Von Bertalanffy, 1938), and a non-linear first-order ODE we call the Canham growth function that is based on a log-normal distribution function (Canham et al., 2004). While we focus on organisms growing, the underlying method is more generally applicable to repeated observation data governed by other dynamics such as smooth motion, and the package is intended to serve as a demonstration for how such models can be implemented in Stan (Stan Development Team, 2024).

From a structural perspective, hmde is an interface to Stan (via RStan (Stan Development Team, 2019)) that provides a set of pre-built Stan models and soadditional functionality to make analysis easier for the end user. Figure 1 demonstrates the workflow starting from longitudinal data where individuals differ in their behaviour over time. By fitting a suitable ODE we can extract individual parameters that allow us to fit functions to each organism, and hence estimate a sequence of sizes over time that accounts for individual variation. O’Brien et al. (2024) showed that fitting individual ODEs in this fashion was able to smooth out measurement error and provide better estimates of the true sizes over time.

**Figure 1:**
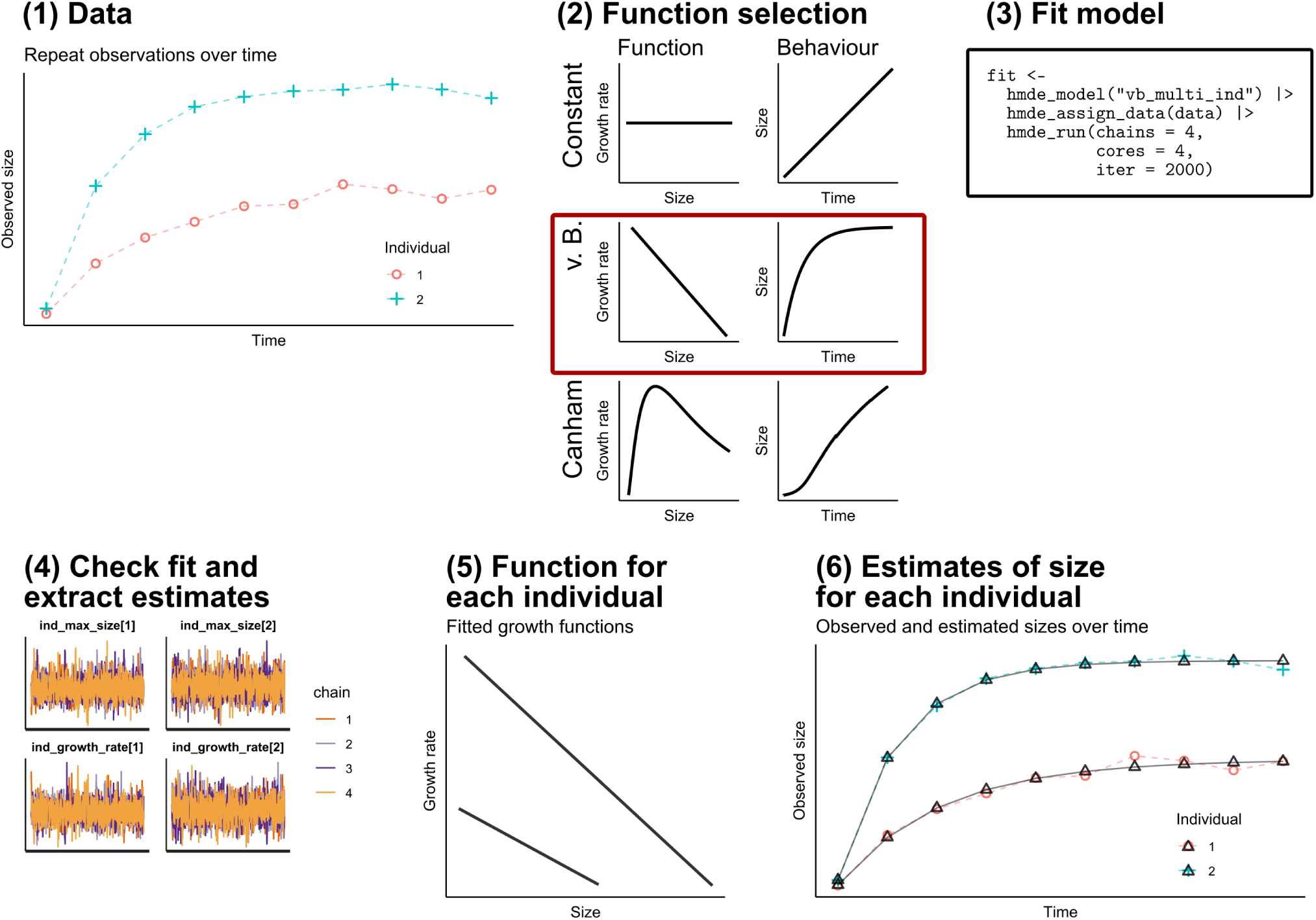
The process of using hmde starts with repeated size observations (1). The user then chooses the most appropriate function from the three available (2), fits the hierarchical model (3), and verifies that the fit converges sufficiently (4). The user then extracts function parameters for each individual, which parameterise the chosen growth function at the individual level (5). As a result of an individual-level growth function, the user also has access to individual size trajectories (6).

In comparison to existing tools, hmde is less flexible but more user friendly than building Stan models directly, while brms (**?**) requires a lot of set-up work to program in the ODE function. Direct ODE solvers such as deSolve (**?**) do not give access to the estimation side.

This paper details the theoretical structure of the underlying mathematics and statistics. We go through the required data structures and include example data to show how the repeat observation structure is represented in the computer. Finally, we provide demonstrations of each implemented model using provided example data, and outline some known statistical issues that may be encountered in user-developed models that rely on numerical integration.

## 1 The Mathematics

In this section we describe the underlying mathematical model that hmde implements, introduce the provided functions, and give some guidance for how a user can determine if, or which, of the provided functions are suitable for their data. We predominantly use size and growth terminology, as we are coming from the perspective of ecology applications where we study growth of individuals over time. The underlying statistical method is however more general. The approach and package could be applied to any ODE problem, where we are trying to parameterise the available ODEs based on observations of the output.

### 1.1 Longitudinal model

We are interested in a size *Y* (*t*) which is governed by the differential equation for growth

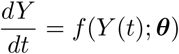

and we take measurements at a sequence of times *t*_1_*, . . ., t_N_*, whose true (error-free) values satisfy

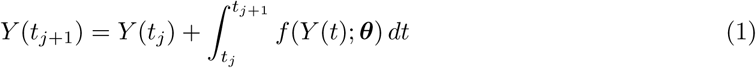

for some vector of parameters ***θ***. We assume that *f* is known (or at least chosen), but the parameter vector needs to be estimated and is the focus of the model.

### 1.2 Pre-built functions

We have implemented three models that are available for direct use with hmde: constant, von Bertalanffy, and Canham. These were chosen to demonstrate a range of implementations across biology with varying data requirements and constraints. The models have one, two, and three parameters respectively, and demonstrate linear and non-linear dynamics. All of the DEs are time independent, in the sense that the time variable *t* does not appear on its own in the DE itself. The von Bertalanffy and Canham models depend on *Y* (*t*) explicitly, which is referred to as size-dependent growth in the ecology literature.

#### Constant

The constant model is given by

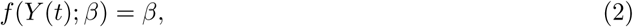

with *β* as the growth rate parameter. The constant model is mathematically equivalent to a linear model for sizes over time. Despite their simplicity, constant models are used, typically in the form of mixed effects models fitting average growth to individuals such as Bhandari et al. (2021) and Lussetti et al. (2019).

#### von Bertalanffy

The von Bertalanffy model (Von Bertalanffy, 1938; Shine and Charnov, 1992) is given by

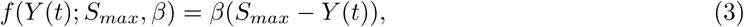

where *S_max_* is the asymptotic maximum size and *β* controls the growth rate. The integral solution to Equation (3) is also known as the monomolecular function, and used in forestry. The implemented version in the pre-compiled model is translated by a sample mean *y* in order to provide more robust and efficient estimation. Translation does not affect the function behaviour as there is a back-transformation for the output estimates.

#### Canham

The Canham function, developed in Canham et al. (2004), specifies a unimodal hump-shaped function for the derivative, given by

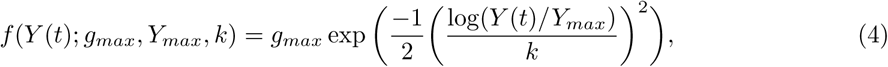

with *g_max_*being the maximum growth rate, *Y_max_* the *Y* -value at which the peak occurs, and *k* a spread parameter that controls how narrow or spread out the peak is.

### 1.3 Choosing a function

There are mathematical considerations for whether any of the existing models will work for a given data set. The constant and Canham models are constrained to have non-negative growth and Canham is not defined for *Y* (*t*) ≤ 0. From a biological perspective non-negative growth may not always be true, for example an organism can reduce in size or mass particularly in the short term. In the constant function case this can be taken as an averaging constraint, while for Canham, the long time between observations of the trees in the demonstration data make non-negative growth a more reasonable assumption as we are smoothing out the behaviour in the five year period. The von Bertalanffy model can fit negative growth and negative *Y* (*t*) even if the latter is biologically impossible. The typical use case for the von Bertalanffy model is for growth that declines as an organism approaches a maximum size, but if the data shows a value that is shrinking asymptotically to a value of *Y* (*t*) (negative growth), the model can still be fit.

The key data requirement is longitudinal data. The point of the model is to fit a function to the dynamics for each sampling unit (individual animal or plant in the data for this paper). The user needs to have repeated observations *y_ij_* from that unit to do so, and know the time of each observation as well. Datasets of pairwise difference data are not suitable, the raw values should be used instead. Consistent identification of individuals is also important, but that is a matter of data cleaning and quality as we have not implemented a structure that allows for misclassified individual.

The choice of DE depends on both data quantity and desired behaviour. From the data quantity perspective, fitting individual-level parameters requires a certain minimum number of observations for each individual, which varies across DEs. The constant function is the most generally applicable, requiring a minimum of two observations to estimate the single function parameter. The von Bertalanffy model has two parameters and a minimum of three observations. Canham, with three parameters, can theoretically be fit to four observations but we strongly recommend at least five. If the chosen function is a good representation of the underlying dynamics, all the models will perform better with more observations per individual.

In terms of desired behaviour, the user should choose a DE with a functional shape that captures the behaviours of interest. The constant model fits an average rate of change to each individual, smoothing out all dynamics. Such a model is unlikely to be realistic across a lifetime, but may be good enough if what is required is a distribution of average growth rates, or if growth appears steady across a time period. The von Bertalanffy model assumes high growth at the smallest sizes, and an asymptotic maximum value which may not be observed in the data. The Canham model has accelerating growth at small sizes, and declining growth after the peak at (*Y_max_, g_max_*). As with the von Bertalanffy model, the decline to asymptotically 0 growth may not be observed in data, as seen in O’Brien et al. (2024).

### 1.4 Numerical methods and analytic solutions

Equation (1) invokes a differential equation that governs the dynamics of *Y* (*t*) which needs to be solved to find *Y* (*t*). We have implemented different methods to solve for *Y* (*t*), depending on the specific DE involved. For the constant growth model hmde uses the analytic solution which is given by

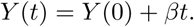

For the von Bertalanffy model hmde uses the analytic solution

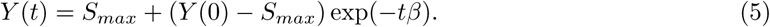

The Canham function in Equation (4) is extremely non-linear and the closed form solution requires the complex inverse error function which is not available in Stan. As such, hmde must employ numerical methods. As Stan provides some ODE solvers, we use the inbuilt Runge-Kutta 4-5 solver for Canham which has an adaptive step size.

In the process of building hmde we encountered a very bimodal posterior distribution arising from error in numerical integration of the von Bertalanffy model. O’Brien et al. (2025) is an extensive investigation of the problem. In hmde itself, we avoid these issues by using the analytic solution of the constant and von Bertalanffy models. For Canham, hmde still uses the RK45 method which O’Brien et al. (2025) found to give negligably biased and unimodal results in simulation.

## 2 The Statistics

The underlying statistical approach in hmde is a hierarchical Bayesian method for inverse problems: attempting to estimate parameters for a DE based on a chosen structure for known statistical relationships governing the distribution of those parameters, and observed data of the resulting process. We assume that the dynamics are continuous, but observations are discrete and finite.^1^

We have implemented two sets of models, the first if a single individual is included in the data, and the second if multiple individuals are, which adds a (hierarchical) cross-individual distribution for parameters. The underlying structure is the same with one additional level for multi-individual data. We have not implemented a multi-population model due to the computational constraints associated with doing so, and because in application to cross-species variation analysis we want to avoid shrinkage towards the mean that a multi-population model would encourage. Aside from an individual effect there is currently no option for covariate data in hmde but this is planned for future releases.

From the ‘bottom’ up, the multi-individual model has the following levels:

### Measurement

The data is repeated measurements on *M* individuals. For the *i*th individual, there are *N_i_*measurements at times *t*_1_*, . . ., t_N__i_*, and we denote the *j*th such measurement as *y_i,j_*. We assume the observation is centred on the true value *Y_i_*(*t_j_*) but is measured with error:

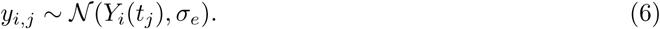

For the Bayesian model, hmde has the measurement-level prior

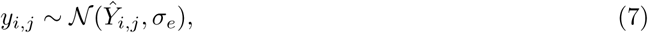

where *Y*^^^*_i,j_* is the estimate of *Y_i,j_* obtained by integrating the growth function for *j >* 1, or the initial condition (modelled as a latent variable) for the first size. O’Brien et al. (2024) demonstrated that a normality assumption works well in the case of a specific normal mixture distribution error process (which is still symmetric and centred at 0), so we are comfortable with that structure. In theory more general error models can be implemented in Stan, but that is best implemented purpose-built for a specific use case.

### Individual

The vector of parameters for *f* is fitted at the individual level, giving ***θ****_i_* as the parameter vector estimate for individual *i*. The form of *f* is fixed for a given model so the individual variation is encoded by different parameter values. For example in the constant model, *β_i_* is the parameter for individual *i*.

### Population

Each parameter in the vector ***θ****_i_* comes from a distribution that operates at the population level. The chosen growth functions are parameterised to use log-Normal priors on the population-level distribution. Consider the growth parameter *β* from the constant model for example, the prior is

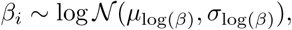

with population-level hyper-parameters that govern the mean and standard deviation of the log-normal distribution. The relationships between individuals within the population are entirely encoded by the parameter distributions.

### Global

The hyper-parameters have their own priors, which are treated as independent across different parameters. hmde uses a normal distribution for the means, and a half-Cauchy distribution for the standard deviation parameters:

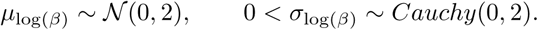

Most means have a chosen default centre value, except for *µ*_log(_*_Smax_*_)_ in the von Bertalanffy model, which instead is centred at the maximum of the log-transformed *y_i,j_*observations. The parameter *S_max_* is the asymptotic maximum size for an individual, so it is reasonable to expect the mean of that distribution to be closer to the maximum observed size than another specified value.

The *σ_e_*error term from Equation (6) also has a prior at the global level given by

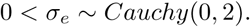

Depending on the use case (*e.g.* working with millimetres rather than centimetres) these priors may be more or less informative. The user is able to change the parameter values but not the underlying distributions by providing the necessary arguments to the hmde assign data function. We demonstrate this functionality later in the paper.

If the user is fitting a model to a single individual, they drop the global level hyper-parameters and fit the individual parameters to a log-normal distribution with specified mean and standard deviation values, for example *β* ∼ log N (0.1, 1) for the constant model with a single individual. The *σ_e_*distribution is preserved in order to estimate the error behaviour.

### 2.1 Estimation

hmde uses a Markov Chain Monte Carlo estimation process, which takes samples from the posterior distribution (Gelman et al., 2021), implemented with a Hamiltonian Monte Carlo algorithm (Stan Development Team, 2024). For each estimator the mean of the posterior samples is the most common estimate (Gelman et al., 2021) and is provided, but hmde also calculates the posterior median and a central 95% credible interval of samples for individual, population, and error parameters via the hmde extract estimates function. The user is able to extract their own estimates from the samples as the hmde run function returns a Stan fit object that includes the sample chains themselves. For some applications the posterior distribution represented by the samples is of interest as well.

## 3 The Data

This section will detail the requirements for using user data with the hmde package and introduce the three demonstration datasets that come with the package.

### 3.1 Data structure

The heart of the statistical model hmde implements is a longitudinal structure for repeated observations over time. The package requires this structure in datasets, with at least two measurements per observational unit at different times, a record of when the measurements were taken to calculate the observation interval, and if there are multiple individuals then a way of identifying which individual each measurement comes from. The basic form is a table – usually a data frame or tibble (Müller and Wickham, 2024) – with columns for observations *y_i,j_*, the observation index *j* counted for each individual, time *t_i,j_*, and individual index *i*. In the following example data, *i* is represented as ind id and *j* as obs index which are the variable names hmde uses internally:

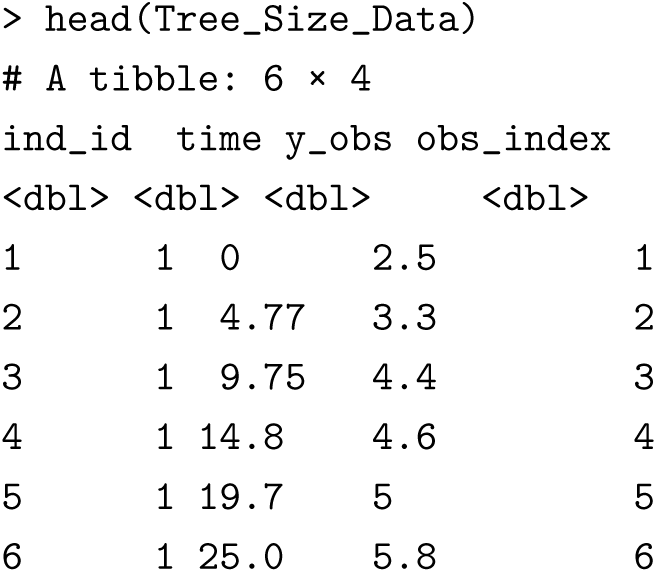

There is a way to pass in data with different names which will be addressed in the section on workflow.

The implemented models have some additional requirements that are typically calculated from the basic framework. For models fitted to single individuals the core structure within the code is

- y obs: a vector of real numbers of length n obs which is the observations,
- n obs: an integer giving the total number of observations which is automatically calculated from the length of y obs,
- obs index: a vector of integers of length n obs giving the index *j* of observations.
- time: a vector of real numbers of length n obs giving the time since the first observation.
- y bar: a real number used for the von Bertalanffy model to centralise the data, typically the mean of the observed values. Automatically calculated from y obs.

For the multi-individual model there are additional values:

- ind id: vector of integers of length n obs that gives the individual index *i*.
- n ind: integer giving the number of individuals in the sample which is automatically calculated from the number of unique values in ind id.

The hmde model() function run on the name of a model will return a hmde model template class object, which is a data structure comprised of the ordered list of numbers, vectors, and strings required for the corresponding Stan model. We have implemented plot, summary, and print S3 standard functions for the hmde model template class, the first of which provides a generic plot of the DE function, using the mean prior parameters, while summary and print return the model name and content of the object.

Here’s the multi-individual Canham template which shows the default priors for the top-level parameters:

­ hmde_model("canham_multi_ind")

$n_obs NULL

$n_ind NULL

$y_obs NULL

$obs_index NULL

$time NULL

$ind_id NULL

$prior_pars_pop_log_max_growth_mean

[1] 0 2

$prior_pars_pop_log_max_growth_sd

[1] 0 2

$prior_pars_pop_log_size_at_max_growth_mean

[1] 0 2

$prior_pars_pop_log_size_at_max_growth_sd

[1] 0 2

$prior_pars_pop_log_k_mean

[1] 0 2

$prior_pars_pop_log_k_sd

[1] 0 2

$prior_pars_global_error_sigma

[1] 0 2

$model

1. [1] "canham_multi_ind"

attr(,"class")

1. [1] "hmde_model_template"

Some of the variable names are very long, because they aim to represent the variable role. For example prior pars pop log size at max growth mean is the vector of prior distribution parameters for the population-level hyper-parameter that controls the mean of the log-transformed *S_max_*(size at max growth) distribution. These typically only come into play when specifically controlling priors, or working with the hyper-parameter distributions.

### 3.2 Provided datasets

For demonstration purposes we have included four in-built datasets that are prepared for immediate use with the existing models. These data represent a range of taxa, both experimental and observational data, and are taken as prepared subsets of public datasets with permission from the original authors.

#### Trout size data

The trout size dataset is taken from the SUSTAIN trout data (Moe et al., 2020), a set of mark-recapture data for *Salmo trutta* in a land-locked population in Norway. We have taken a stratified sample of 50 individuals, where the strata are the number of observations from the individual fish:

- 25 individuals with 2 observations,
- 15 individuals with 3 observations,
- 10 individuals with 4 observations.

Within strata, the individuals were selected by a simple random sample without replacement from a sampling frame of individual IDs. The size of the trout is measured in centimetres from end to end. As the survey structure for the trout data requires re-capture, there is no control on the time between observations, which is measured in years. Due to the limitation on observations we have chosen the trout survey as demonstration data for the constant function, which is not size-dependent and has only a single parameter (the average rate of change) to estimate.

#### Lizard size data

Our example lizard data comes from experimental data used in Kar et al. (2024) from the species *Lampropholis delicata*. Size-structured growth based on a von Bertalanffy model has been used in the literature for other lizard and reptile species (Ramírez-Bautista et al., 2016; Shine and Charnov, 1992), so we considered this data a good match for that function. Measurements are in millimetres from the lizard’s snout to the top of the cloacal opening (snout-vent-length or SVL). Time is measured in days. We took a simple random sample without replacement of 50 individuals using the individual IDs as a sampling frame.

#### Tree size data

The tree size data comes from Barro Colorado Island (Condit et al., 2019), in this case from the species *Garcinia recondita*. A simple random sample of 50 individuals was taken from the 400 *G. recondita* individuals used in O’Brien et al. (2024), which were already processed through data cleaning and filtered to have 6 observations each (i.e. have survived the 25 years of observation) and checked for preservation of stem and tree IDs over time. Further filtration chose individuals with more than 3 cm observed difference between first and last size to avoid model fitting problems with the smaller dataset. Size is given as diameter at breast height (DBH) in centimetres, and time is measured in years.

### Additional functions

We have provided some additional functions to help with model diagnostics and analysis.

#### Estimate extraction

hmde extract estimates takes as arguments the fitted model, and a data frame or tibble comprising the y obs, obs index, time, and ind id data. The function returns a list of estimated values at the measurement, individual, and population level (if the model was fit to multi-species data).

#### Fit diagnostics

We provide two additional functions for use alongside standard RStan diagnostics:

hmde extract Rhat takes as argument a fitted model and returns a named vector of *R*^^^ values, excluding those associated with the constant generated quantities used to confirm prior parameters. The constant generated quantites produce NAs as *R*^^^ values because they have no variance.

hmde plot Rhat hist takes as argument a fitted model and returns a histogram of *R*^^^ values that excludes those associated with the constant generated quantities.

#### Model plotting

There are two purpose-built functions to plot the fitted model:

hmde plot de pieces requires the output of hmde extract estimates, with optional arguments to change the *x* and *y* axis labels, colour, and opacity of the plotted lines. Returns a plot that displays each individual’s fitted DE, from the first to last estimated size.

hmde plot obs est inds requires the output of hmde extract estimates, a tibble or data frame of the measurement data used to fit the model, and an integer for sample size or a vector of ID values matched to the data. Optional arguments change the *x* and *y* axes and title. Returns a plot of observed and estimated size over time for either a chosen number of them randomly selected from the data, or specified individuals based on the vector of ID values.

## 4 The Workflow

All of the models provided in hmde leverage the same workflow. In the next sections we walk through implementation but first we detail the workflow in theory. As an example, let’s say we choose the constant function as the model. The following code will take the provided trout size data, convert it in to the structure required to fit a constant model, fit the model, and extract the posterior estimates at measurement, individual, and population levels. We store the model fit output from step 4 in order to save the fit for traceplots or other diagnostic analysis.

\# Constant function chosen trout_constant_fit <- hmde_model("constant_multi_ind") |> hmde_assign_data(data = Trout_Size_Data) |> hmde_run(chains = 4, cores = 4, iter = 2000) trout_estimates <- hmde_extract_estimates(trout_constant_fit, input_measurement_data = Trout_Size_Data)

The hmde assign data can take different configurations of data. If passed a data frame with column names y obs, obs index, time, and ind id in any order it will use those columns directly. Data can be individually assigned to those names as arguments, for example

hmde_model("constant_multi_ind") |> hmde_assign_data(y_obs = Trout_Size_Data$y_obs, obs_index = Trout_Size_Data$obs_index,

time = Trout_Size_Data$time, ind_id = Trout_Size_Data$ind_id)

will produce the same data list as

hmde_model("constant_multi_ind") |> hmde_assign_data(data = Trout_Size_Data)

and can be used for data frames with different column names. There are some internal checks for data size consistency to help with passing in individual vectors.

Returning to the fitted model, here are the first few rows of each element in the estimates list:

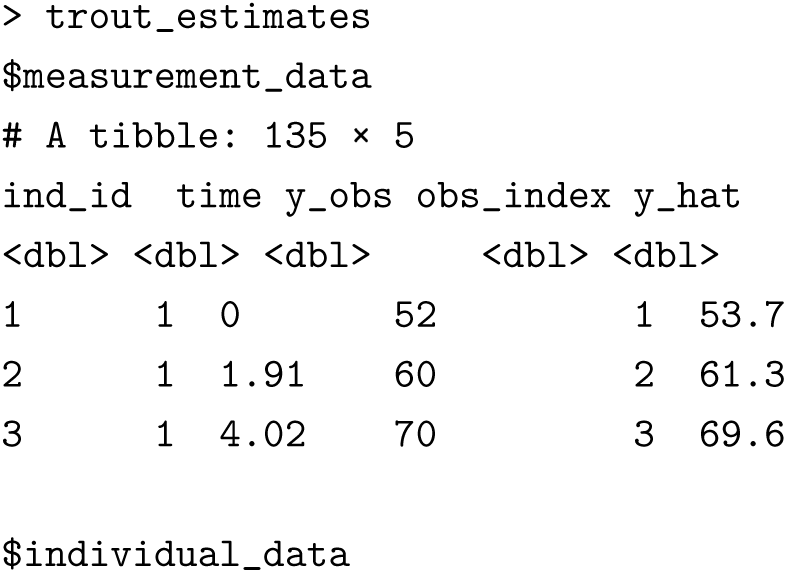
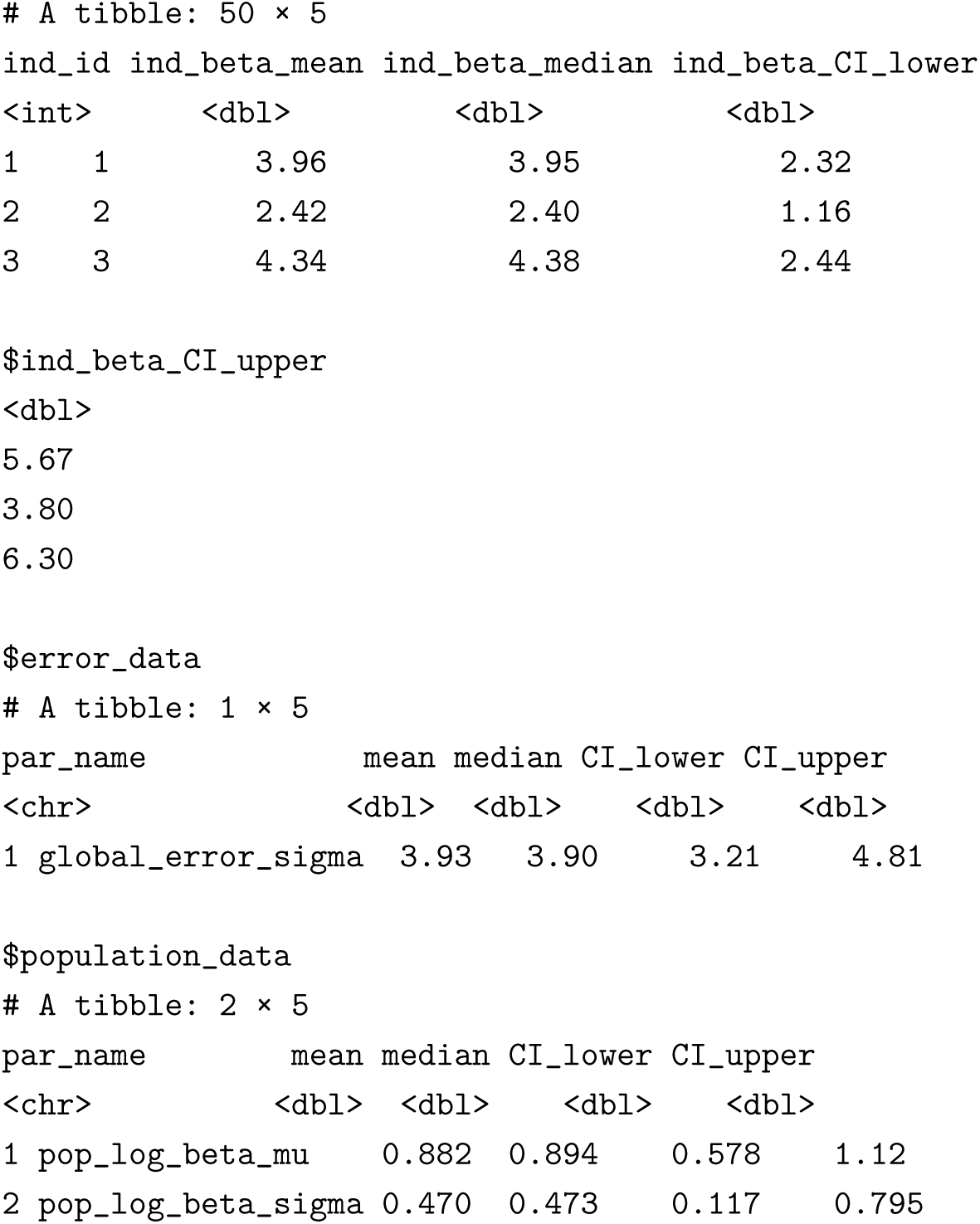

Each tibble can be extracted from the list and used as a regular tibble/data frame structure for further analysis. Some examples include comparisons of observed and estimated sizes using the measurement data, parameter scatter plots and histograms from the individual data estimates, and distribution information encoded in the hyper-parameters in population data values. Examples of analyses are given below for the case studies we use, and relevant code can be found in the vignettes.

In greater detail, the 5 steps to the fundamental workflow for hmde are:

1. **Choose model:** To see a list of model names run hmde model names().
2. **Data structures:** hmde requires specific data structures for the Stan models. A detailed list of each can be seen by running the function hmde model and giving it the name of the relevant model. The user can build each element in the list themselves, or construct a table with observations from each individual, an ordering of observations for the individual, the time at which they occurred, and information on which individual the observation came from. The table structure of ind id, time, y obs, and obs index is the easiest to work with as hmde assign data is built to use those column names and the list output by hmde model to give the Stan data structure.
3. **Convert data to** Stan **model structure:** The Stan model requires a specific list structure with agreement between the lengths of some vectors and corresponding integer values. Hmde provides the hmde assign data function that converts provided data into the required list format and checks for the necessary size agreements. There are a handful of ways to pass datasets depending on the level of control required. A data frame or tibble structure with columns named y obs, obs index, time, and ind id can be passed as the data argument and hmde assign data will format the list for the model. Specific list elements can be assigned directly by passing an argument of that name to hmde assign data, with the value of the argument being the data to be assigned.
4. **Run the model:** The function hmde run loads the chosen model and runs the Stan MCMC sampler on the provided data, returning a Stan fit object. If no control parameters are provided the sampler will run 4 chains with 2000 iterations on a single CPU core as default for Stan. Sampler controls can be passed to hmde run as they would to the sampler directly, for example chains = 2 will run only two chains.
5. **Extract posterior estimates:** hmde has hmde extract estimates to make posterior pro- cessing easier. The function takes the Stan fit object output by hmde run and the data it was fit to in the tibble structure described in step 2. Which model was chosen is named in the Stan fit object. The output is a list of tibbles with posterior parameter estimates for each level in the hierarchical structure. The function gives the mean of posterior samples as the estimate for *Y*^^^*_i,j_*, but additionally the posterior sample median, and a central 95% credible interval for the individual- and population-level parameter estimates.

Steps can be strung together using the pipe operator |> (native to R, the Magrittr pipe %>% Bache and Wickham, 2022 also works) which uses the output of a previous function as the **first** argument of the next.^2^ We recommend retaining the output Stan fit object from hmde run for diagnostic purposes. The following demonstrations align with the vignettes for each model, and demonstrate each included DE and associated provided dataset. We go through the constant model in detail, summarise the most interesting results from the von Bertalanffy model, and give a brief overview of the Canham results, which are more extensively explored in O’Brien et al. (2024). Due to the random nature of MCMC sampling, we have used the set.seed function to provide replicable results.

### 4.1 Constant growth fit to trout size data

In circumstances where the number of observations available per individual is very limited, average growth rates over time may be the only plausible model to fit. In particular, if there are individuals with only two size observations the best that can be done is a single estimate of growth rate based on that interval. Such a model behaves as constant growth, which we can think of as the average rate of change across the observation period and is given by Equation (2), *β* is the average growth rate across the observation period. The constant growth model corresponds to linear sizes over time, and is equivalent to a linear mixed model for size, where there is an individual effect when fit to multiple individuals.

Example data for the constant model comes from Moe et al. (2020), a publicly available dataset of mark-recapture data for *Salmo trutta* in Norway. The time between observations is not controlled, nor is the number of observations per individual. As a result the data consists primarily of individuals with two observations of size, constituting a single observation of growth which limits the functions that can be fit to individuals, as a single parameter model is the best that can be fitted to two sizes. The constant growth function in Equation (2) is the most appropriate of the functions in hmde, where the single growth interval is used to estimate the average growth rate *β*.

To implement the workflow, we fit the model and extract the estimates. We have already chosen the constant model for step 1, so to look at the required data structure we call hmde model("constant multi ind").

As the provided trout data is already in the form required by hmde assign data we do not need to do any further pre-processing for step 2 and can pass the data directly to step 3 using hmde assign data("constant multi in data = Trout Size Data). The following code includes command to run with multiple cores and demonstrates the use of the pipe operator to pass the required data structure from hmde model to hmde assign data, then the correctly formatted list from hmde assign data to hmde run for step 4 that fits the model. The overall output of this part of the workflow is the model fit.

\# Constant model chosen as Step 1 trout_constant_fit <- hmde_model("constant_multi_ind") |> #Step 2 hmde_assign_data(data = Trout_Size_Data) |> #Step 3 hmde_run(chains = 4, cores = 4, iter = 2000) #Step 4

Fitting the model is the most computationally expensive part, so we recommend storing the model fit at this point.

If the user wants to change the default prior values, add the required prior parameter values as arguments to hmde assign data, which then get passed to the Stan model. The following shows how to change the parameters for the population-level prior on *µ*_log(_*_β_*_)_ to be a mean of 1, and a standard deviation of 3:

#Look at the default prior values hmde_model("constant_multi_ind")$prior_pars_pop_log_beta_mu

#Assign new value

data <- hmde_model("constant_multi_ind") |> hmde_assign_data(data = Trout_Size_Data, prior_pars_pop_log_beta_mu = c(1, 3))

Assignment can be checked by looking at the value in the template before the model is run:

­ data$prior_pars_pop_log_beta_mu [1] 1 3

There are generated values in the model fit object that return the used prior parameters, (the names all start with check prior pars) so users can double check that the priors they want are actually used.

In step 5 we extract estimates for each level in the hierarchy using the function hmde extract estimates, which is set up to take the data as structured for the model fit.

trout_estimates <- hmde_extract_estimates(#Step 5 fit = trout_constant_fit,

input_measurement_data = Trout_Size_Data)

We directly compare the observed sizes over time to estimated values. For a quantitative fit metric we use *R*^2^ calculated on (*y_ij_, Y*^^^*_ij_*), and root mean squared error (RMSE) calculated as

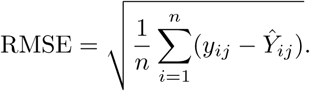

For qualitative analysis we look at scatter plots of observed and estimated sizes, and inspect plots of sizes over time, as in Figure 2(a) and (b). The *R*^2^ statistic is a metric primarily used in linear regression that measures the proportion (ie. decimal value in the [0,1] interval) of variance in one coordinate that can be explained by the regression model. In this context, we interpret it as how strongly the fitted and observed values agree. As statisticians we never expect perfect agreement between a model and data, so we expect *R*^2^ *<* 1, but as *R*^2^ is susceptible to extreme values we use RMSE as a second goodness of fit metric. O’Brien et al. (2024) showed that the change between observed and fitted values can actually correct for measurement errors in size, so disagreement is not a bad thing overall. In this case, *R*^2^ = 0.953 and *RMSE* = 2.75, which indicates strong agreement between the estimated and observed sizes, even though we have chosen a very simplistic model in the constant function.

**Figure 2:**
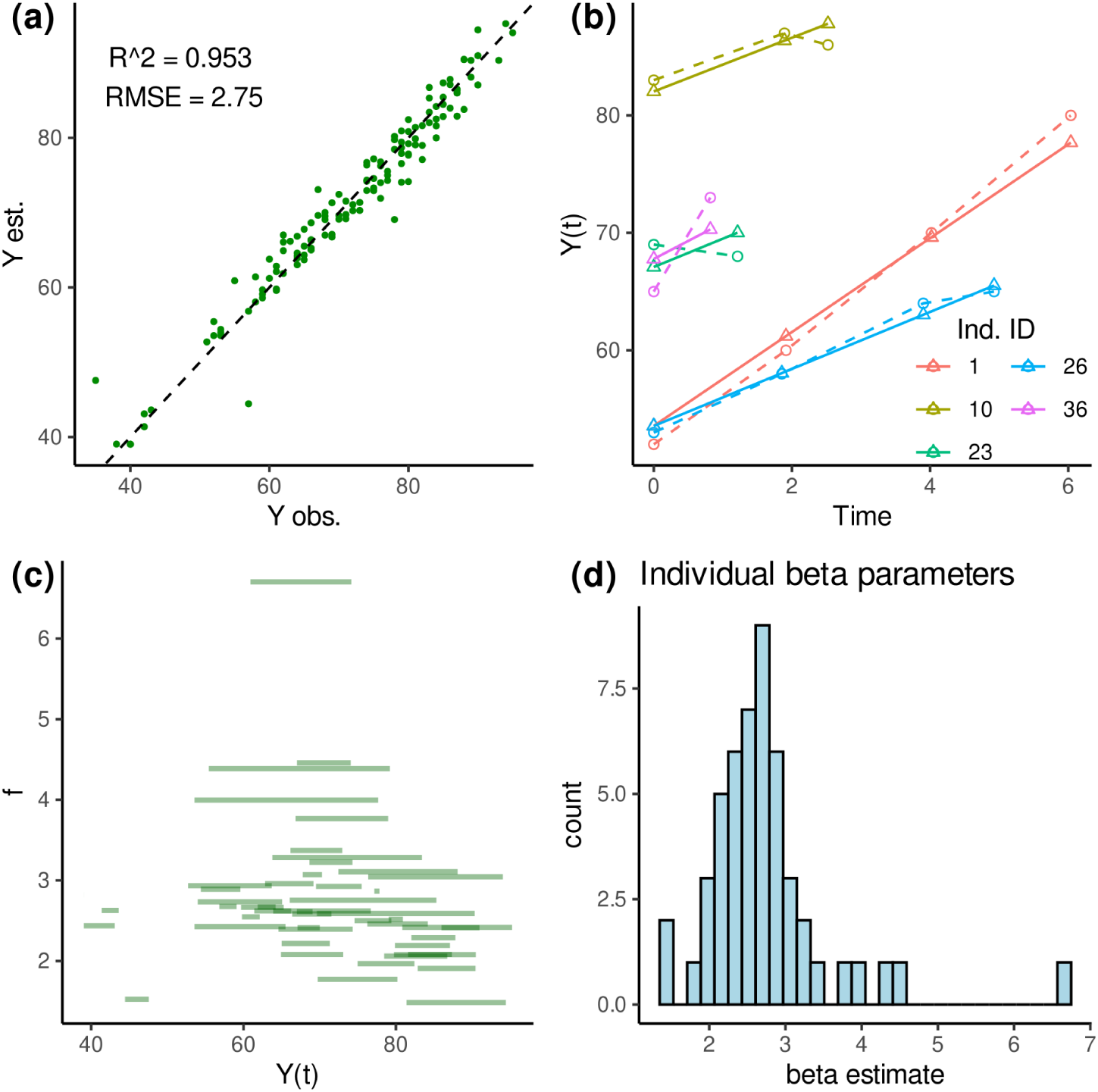
Analysis plots for *S. trutta*. **(a)** gives a comparison of estimated and observed sizes showing strong agreement along the (dashed) line of identity. **(b)** gives observed and estimated sizes over time for five randomly selected indi- viduals using the hmde plot obs est inds function. **(c)** shows all fitted growth functions and was produced with the hmde plot de pieces function. **(d)** is a histogram of the *β*^^^*i*s.

Figure 2(b) demonstrates that at the individual level, the constant growth function produces linear sizes over time that are averaging out the observed behaviour. This plot can be produced from the hmde function hmde plot obs est inds using the measurement data tibble:

hmde_plot_obs_est_inds(n_ind_to_plot = 5, measurement_data = trout_estimates$measurement_data)

Panel (c), a plot of growth functions fit to each individual in the sample, can also be easily produced using the function hmde plot de pieces using the extracted estimates:

hmde_plot_de_pieces(trout_estimates)

In Panel (d) there is a large range of estimated average growth rates, and a possible downwards trend for large sizes that would be better estimated with a size-dependent model.

Finally, analysis of the distribution of *β_i_*, which is done through a histogram, and extraction of the species-level parameter estimates. Figure 2(d) shows a right-skewed, unimodal distribution of average growth rates with a tail of high values. There is one extreme growth value (about 7 cm per year) which may not be biologically plausible, and that individual could be further investigated. Table 1 gives the posterior estimates for the hyper-parameters, with both the raw mean for the log-transformed distribution, and the exponent of that value which can be more easily interpreted as an average growth rate for the species in cm/yr.

**Table 1:**
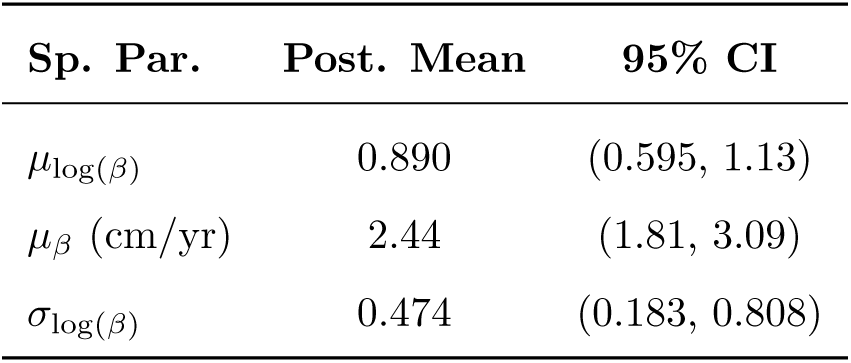
Posterior estimates for species-level hyper-parameters in the constant model for trout. This data comes from the population data tibble in the extracted estimates.

| Sp. Par. | Post. Mean | 95% CI |
| --- | --- | --- |
| $\mu_{\log(\beta)}$ | 0.890 | (0.595, 1.13) |
| $\mu_{\beta}$ (cm/yr) | 2.44 | (1.81, 3.09) |
| $\sigma_{\log(\beta)}$ | 0.474 | (0.183, 0.808) |

### 4.2 von Bertalanffy fit to lizard size data

Our second example uses size-dependent growth based on the von Bertalanffy function given in Equa- tion (3). The key behaviour of the von Bertalanffy model is a high growth rate at small sizes that declines linearly as the size approaches *S_max_*. This manifests as growth slowing as an individual matures, with an asymptotic final size. hmde restricts *β* and *S_max_* to be positive, and uses the maximum of observed sizes as the prior mean for *S_max_*to avoid model pathologies.

The data for the von Bertalanffy demonstration is the Lizard Size Data object provided with the package. As with the constant model, we pass the Lizard Size Data directly through the workflow to fit the model. The following code covers the steps in the workflow in two concise parts.

\# von Bertalanffy model chosen for Step 1 lizard_fit <-

hmde_model("vb_multi_ind") |> #Step 2 hmde_assign_data(data = Lizard_Size_Data) |> #Step 3 hmde_run(chains = 4, cores = 1, iter = 2000) #Step 4

lizard_estimates <- hmde_extract_estimates(#Step 5 fit = lizard_fit,

input_measurement_data = Lizard_Size_Data)

We use plots of sizes over time and estimated growth functions to get a feel for plausibility based on how well the models fit the data. As the von Bertalanffy model has two individual parameters, the distribution of all individual parameter estimates can be scrutinised in a scatter plot. If the user wishes to test for relationships between parameters across individuals, these estimates allow that to be done. Figure 3 gives plots for the analysis based on the extracted estimates. Panels (b) and (c) are produced with the provided functions hmde plot obs est inds and hmde plot de pieces respectively.

**Figure 3:**
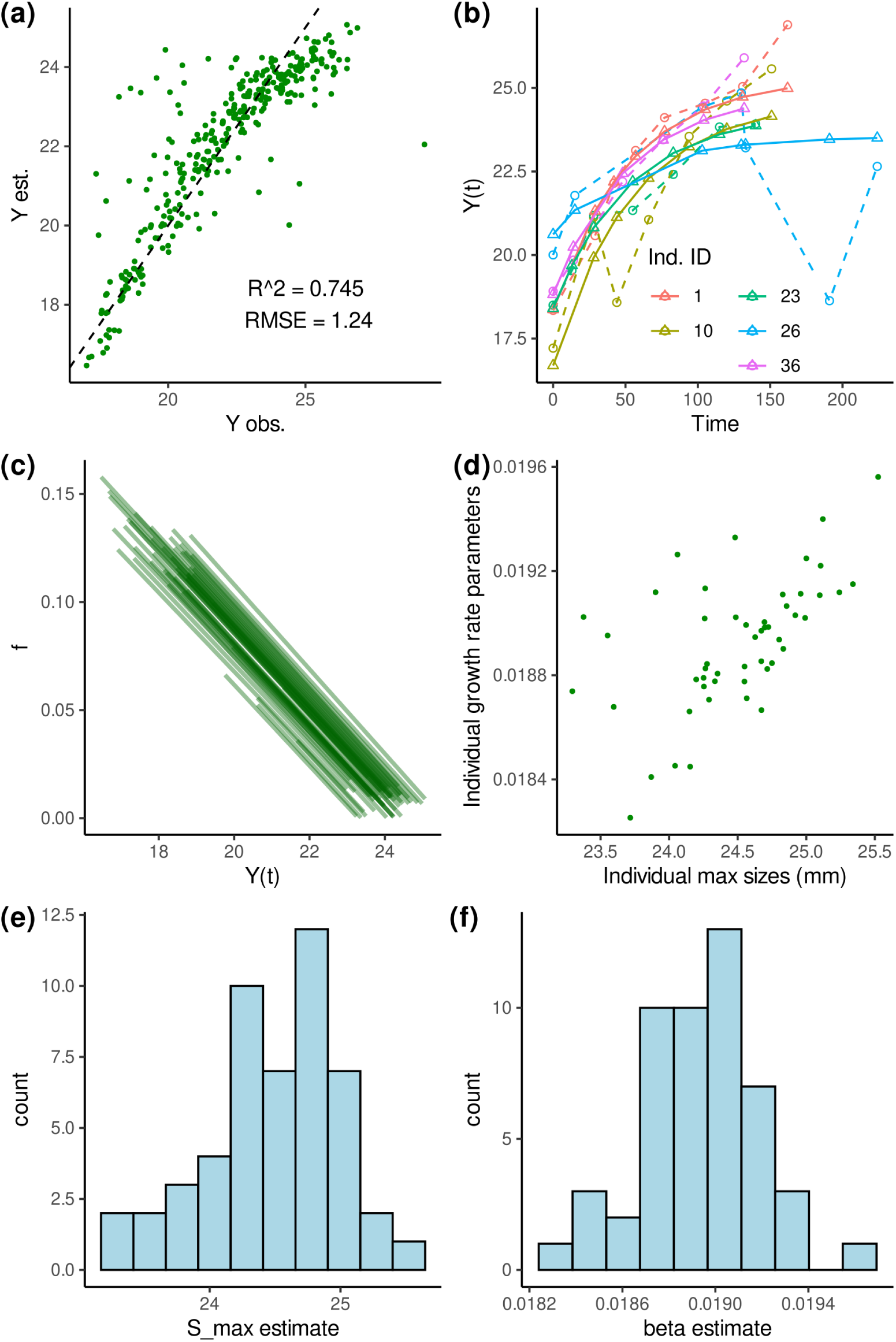
Plots for *L. delicata*. **(a)** gives a plot of observed and estimated *Y* (*t*) values showing reasonable agreement, with some under-estimation of large sizes. **(b)** shows sizes over time for 5 randomly selected individuals. **(c)** gives the fitted von Bertalanffy functions for all individuals and shows that the slopes controlled by *β* are very similar, but there is more variation in the max size. **(d)** is a scatter plot of individual parameters. **(e)** and **(f)** are histograms of individual parameters showing roughly symmetric distributions within the sample.

One interesting result of fitting individual trajectories is that the von Bertalanffy model visibly underestimates the largest sizes in Figure 3(a). The mathematical interpretation of this is that the straight line decay to zero growth may not be appropriate (*i.e.* more growth happens at larger sizes than the von Bertalanffy model predicts), and there may actually be an asymptotic decline in growth instead, necessitating a different growth function for further analysis.

Finally, the population-level parameter estimates. As in the constant growth case, both the raw and exponentiated values for the mean of the log-normal distribution are useful for interpretation, which requires post-processing of the estimates to exponentiate values. The estimates and CIs are provided in Table 2. The estimate of *µ_Y__max_* = 24.5 mm, the average value of maximum size across the population, is quite reasonable given what is known about the species *L. delicata*.

**Table 2:** Posterior estimates for species-level hyper-parameters in the von Bertalanffy model for lizards taken from the lizard estimates$population data tibble. Exponentiated parameters are calculated from the tibble.

| Sp. Par. | Post. Mean | 95% CI |
| --- | --- | --- |
| $\mu_{\log(Y_{max})}$ | 3.198 | (3.176, 3.222) |
| $\mu_{Y_{max}}$ (mm) | 24.48 | (23.95, 25.09) |
| $\sigma_{\log(Y_{max})}$ | 0.0323 | (0.0141, 0.0504) |
| $\mu_{\log(\beta)}$ | -3.979 | (-4.228, -3.762) |
| $\mu_{\beta}$ | 0.0187 | (0.0146, 0.0232) |
| $\sigma_{\log(\beta)}$ | 0.0783 | (0.0112, 0.224) |

### 4.3 Canham fit to tree size data

Our final demonstration implements Equation (4) to model size-dependent growth in trees. We use the Barro Colorado Island data included with the package, which has six observations five years apart for each of 50 individuals from *G. recondita*. The Canham function is used as O’Brien et al. (2024) demonstrated that it performs well for the same data.

The following code runs the Canham model sampling and extracts samples. As sampling takes a few hours for this example we recommend using the provided set of estimates in the Tree Size Ests data object that comes with the package in place of the tree estimates object in this code.

\# Canham function chosen for Step 1 tree_fit <- hmde_model("canham_multi_ind") |> #Step 2

hmde_assign_data(data = Tree_Size_Data) |> #Step 3 hmde_run(chains = 4, cores = 4, iter = 2000) #Step 4

tree_estimates <- hmde_extract_estimates(#Step 5 fit = tree_fit,

input_measurement_data = Tree_Size_Data)

Figure 4 demonstrates a strong alignment between the fitted growth functions and observed growth behaviour, both in Panel (a) with the randomly selected size trajectories and the observed and estimated sizes in Panel (c). Among the individual-level parameter estimates, the Spearman’s rank correlation coefficients were 0.195 for *g_max_*and *S_max_*, -0.511 for *g_max_*and *k*, and -0.257 for *S_max_*and *k*, so there is some evidence of a moderate negative relationship between *g_max_* and *k*, which in practice means that a spike to a higher growth rate is not sustained for a long period, while the lower growth rates with the higher *k* values are more sustained.

**Figure 4:**
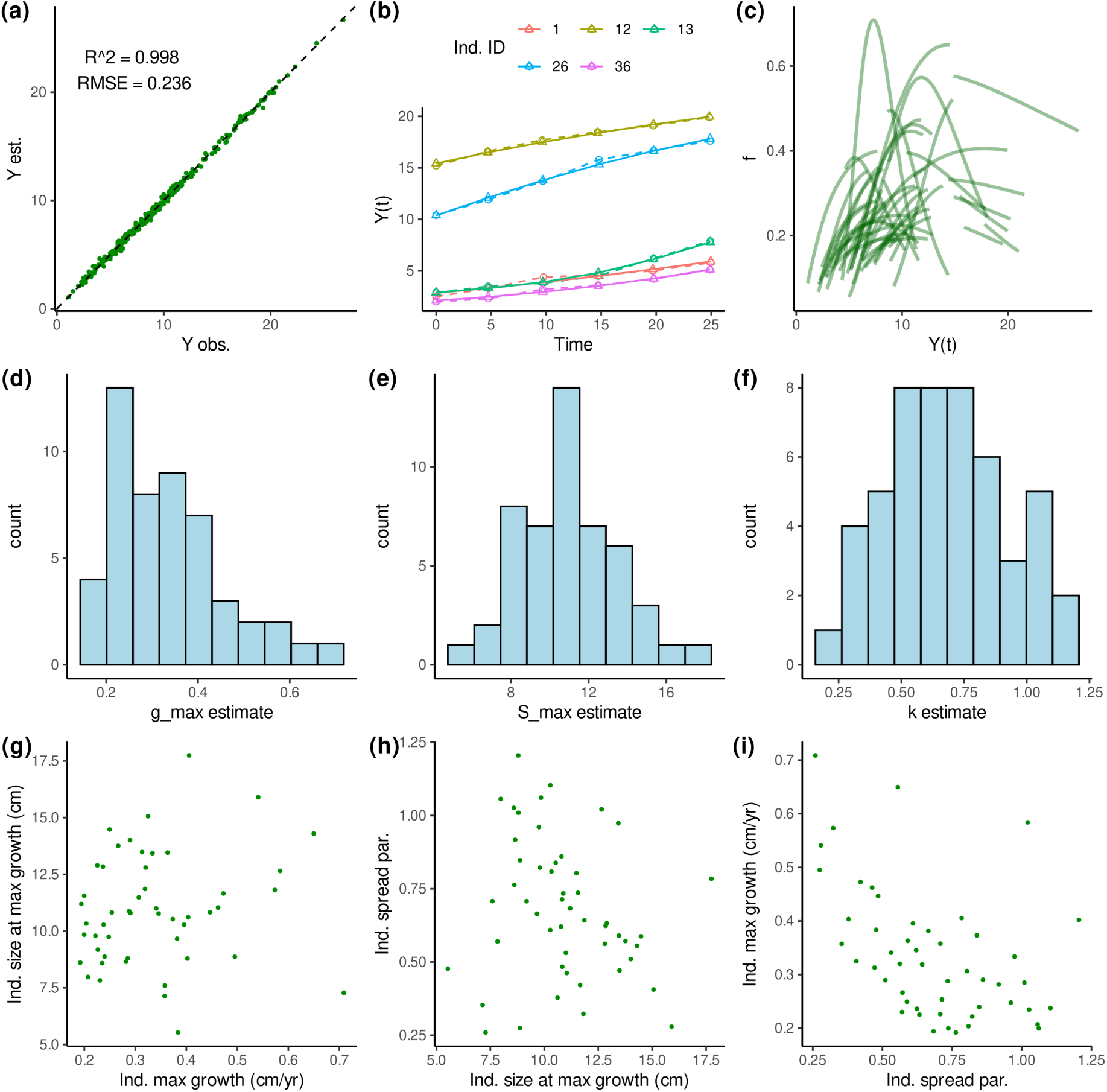
Analysis plots for *G. recondita*. **(a)** is a scatter plot of observed and estimated sizes showing very strong agreement. **(b)** shows sizes over time for 5 randomly selected individuals. **(c)** gives the fitted Canham functions for all individuals. **(d)**, **(e)**, and **(f)** are histograms of individual parameters, with the horizontal axis aligned to the pairwise scatter plots in **(g)**, **(h)**, and **(i)**.

In Table 3 there are higher estimates for the *g_max_* mean parameters compared to the estimates from O’Brien et al. (2024). Bias is to be expected, because we specifically sampled individuals with more than 3 cm difference between the largest and smallest observed sizes as the sampling frame for Tree Size Data, which excludes a lot of very low-growth individuals that would pull down the distribution of *g_max_*. The other parameters show strong agreement to results from O’Brien et al. (2024), with overlap in the 95% CIs.

**Table 3:**
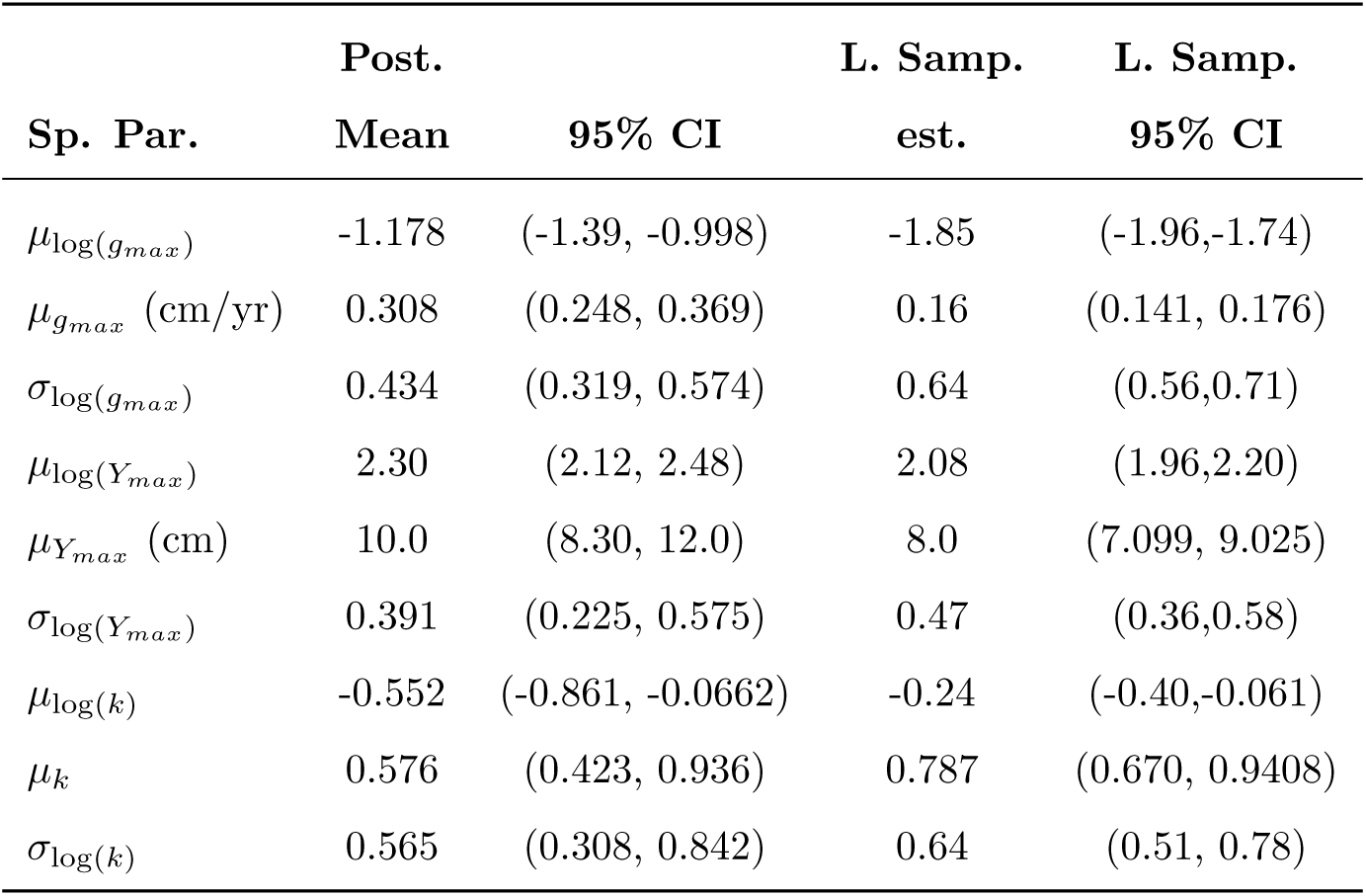
Posterior estimates for species-level hyper-parameters in the Canham model for trees, and the large sample (L. Samp.) estimates from O’Brien et al. (2024) for the same parameters on the larger sample.

## 5 Discussion

hmde provides a baseline implementation of three hierarchical Bayesian longitudinal models in order to demonstrate the application of the method to different biological situations. The package offers an interface to the methods used in O’Brien et al. (2024) which would not otherwise exist. We expand the application to non-tree taxa in this package with two animal demonstrations on *S. trutta* data from Moe et al. (2020) and *L. delicata* data from Kar et al. (2024).

While the method implement in hmde is new, comparisons to existing ones are possible. The constant model is equivalent to a linear model for size as a function of time, with an individual effect. The mathematical expression is

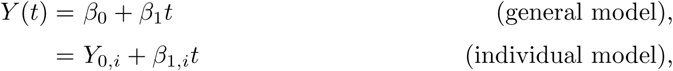

which linear mixed effects and hierarchical linear models can fit such as Lussetti et al. (2019). The explicit inclusion of time enables a longitudinal structure. We nevertheless included the constant model in the package as the simplest use-case with minimal data requirements. Some longitudinal surveys, such as the mark-recapture surveys including the SUSTAIN trout data (Moe et al., 2020), can be dominated by individuals with only two observations which limits the capacity to fit more complex models. If a distribution of average growth rates is what the user wants, the constant model will provide it.

For the von Bertalanffy model, the analytic solution in Equation (5) is exponential and requires a non-linear form of function fitting. Ramírez-Bautista et al. (2016) used a non-linear model fit to pairwise difference data, and fit the von Bertalanffy model at the population level with a sex effect rather than fitting individual effects. In contrast we fit an individual longitudinal model where the population-level behaviour is encoded in the distribution of individual parameters.

The Canham model is the stand-out of the three implemented in hmde, as the absence of an analytic solution to Equation (4) means hmde needs to use numerical integration methods to encode the longitudinal structure. Fitting the solution through the longitudinal model given in Equation (1) contrasts to population-level average trajectory models fit as functions to pair-wise difference data such as Herault et al. (2011). The population model operates at the level of the DE and does not attempt to estimate a size over time trajectory for individuals. I get access to the individual trajectories directly, enabling deeper analysis of behaviour within a population.

Mixed effects models such as Bhandari et al. (2021) uses covariate information to better fit models for tree growth, while Ramírez-Bautista et al. (2016) used a sex effect for lizards. Environmental covariates such as light and spatial dynamics are a key part of ecological models (Falster et al., 2018; Rüger et al., 2022; Babcock et al., 2012). The best that can currently be done in hmde is to run independent models across a single grouping variable. The option to include covariate data is a planned feature for hmde.

By providing demonstrations from three different taxa we show some of the applicability of a very general underlying method for longitudinal data. We chose to implement three models: constant (average) growth, von Bertalanffy (linear first order ODE – linear with translation), and the Canham (Canham et al., 2004) function (non-linear first order ODE). Future work could expand applications to other models, as even within tree growth modeling there are many other functions for dynamic processes (Herault et al., 2011). Another extension would be investigate dynamics in more than one dimension for *Y* such as 3D volume models of size, or two- or three-dimensional motion based on a position-momentum model.

## 6 Conclusions

The hmde package provides a relatively user-friendly structure for implementing the chosen set of models with a hierarchical Bayesian longitudinal method. We have given demonstrations of each model matched to an included dataset of repeat survey or experimental measurements that suit the chosen function.

## Footnotes

1 In the Bayesian inverse methods literature finite observations with measurement error is known as a situation of practical identifiability (Latz, 2023) and presents additional difficulties due to limited information compared to infinite, zero-error data which is used theoretically. More work needs to be done to look at the theoretical side of the model hmde implements.

2 Pipe is perhaps easiest thought of as a way to turn recursive function calls into an easier-to-read chain of them. From a mathematical perspective it is function composition where f(x) |> g() is the same as *g*(*f* (*x*)) = *g ◦ f* (*x*). Thankfully, the arrow points to the next function so the reading order is easier for English native speakers.

